# The Formation of a Stable Sliding Clamp Discriminates MSH2-MSH3 and MSH2-MSH6 Mismatch Interaction

**DOI:** 10.1101/2021.10.21.465318

**Authors:** Brooke M. Britton, James A. London, Juana Martin-Lopez, Nathan D. Jones, Jiaquan Liu, Jong-Bong Lee, Richard Fishel

## Abstract

MutS homologs (MSH) are highly conserved core components of DNA mismatch repair (MMR). Mismatch recognition provokes ATP-binding by MSH proteins that drives a conformational transition from a short-lived lesion-searching clamp to an extremely stable sliding clamp on the DNA. Once on DNA the MSH sliding clamps provide a platform for the assembly of MMR strand-specific excision components beginning with the highly conserved MutL homologs (MLH/PMS). Previous studies with short mismatch-containing oligonucleotides revealed an MSH ATP hydrolysis (ATPase) cycle that included mismatch recognition, the formation of an ATP-bound sliding clamp and dissociation from the end of a mismatched DNA that ultimately recovers the mismatch binding conformation. We found that ATP-bound MSH complexes on blocked-end or very long DNA are extremely stable under a range of ionic conditions. These observations underpinned the development of a high-throughput fluorescence resonance energy transfer (FRET) system capable of clearly distinguishing between HsMSH2-HsMSH3 and HsMSH2-HsMSH6 activities that is suitable for chemical inhibitor screens.

## INTRODUCTION

Mismatch repair (MMR) corrects polymerase misincorporation errors, selected chemical and/or physical DNA damage as well as sequence heterology resulting from DNA recombination (1). Homologs of the prototypical *E.coli* (Ec) EcMutS and EcMutL are the most highly conserved MMR components across biology. The function(s) of the MutS homologs (MSH) and MutL homologs (MLH/PMS) have been of interest for decades. It is well known that MSH proteins recognize mismatched nucleotides (2, 3). However, the operation of these essential components following mismatch recognition has been persistently unsettled (1,4–6).

MSH proteins function as dimers or heterodimers with each half containing a highly conserved Walker A/B nucleotide-binding motif (7). In addition to simple nucleotide mismatches, MSH proteins recognize insertion-deletion loop-type (IDL) mismatches and several relatives have evolved to recognize recombination cross-over structures during meiosis (8–10). The MSH search and recognition process has been shown to involve the formation of an incipient clamp that interrogates the DNA by rotation coupled diffusion (11, 12). ATP binding by all MSH proteins results in the formation of a stable freely diffusing sliding clamp on the DNA (8,10,12–17). The biochemical function of the MLH/PMS proteins had been largely enigmatic until recent single molecule imaging studies showed that the EcMutS sliding clamp provides a platform for EcMutL to form a second extremely stable sliding clamp on DNA containing a mismatch (18). In *E.coli*, these two sliding clamps function together and/or separately to enhance EcMutH DNA association and GATC-hemimethylation incision activity (18–20) as well as EcUvrD strand specific unwinding-displacement activity (21). The human HsMSH2-HsMSH6 and HsMLH1-HsPMS2 similarly form a cascade of sliding clamps, although the detailed downstream functions remain poorly understood (22).

Mutation of HsMSH2, HsMSH6, HsMLH1, and HsPMS2 cause the most common cancer predisposition in humans known as Lynch syndrome or hereditary non-polyposis colorectal cancer (LS/HNPCC) (23, 24). This observation is consistent with a pathogenic dysfunction of the major HsMSH2-HsMSH6 or HsMLH1-HsPMS2 heterodimers. Most cells that contain these major human heterodimers also possess the HsMSH2-HsMSH3 heterodimer (9, 10). HsMSH2-HsMSH3 primarily recognizes large IDL’s, while HsMSH2-HsMSH6 recognizes a broad array of common replication errors that includes single nucleotide mismatches and small IDL’s (10, 25). Mutations of HsMSH3 have not been found to cause LS/HNPCC (26–28). However, mutation of HsMSH3 appears to eliminate the pathogenic expansion of trinucleotide repeat (TNR) disease genes such as Huntington’s disease and Myotonic dystrophy (29). Together, these observations suggest that hampering HsMSH2-HsMSH3 activity, without affecting HsMSH2-HsMSH6 functions, could engender a useful therapeutic for TNR disease pathologies while not encouraging cancer.

Here, we have employed a variety of bulk and high-resolution biophysical methods to dissect MSH progression to a stable sliding clamp. We found that the mismatch binding peaked at relatively low ionic strengths, similar to previous studies (13,16,30). The dissociation kinetics and lifetime of the ATP-bound MSH sliding clamp determined by surface plasmon resonance (SPR) analysis and single molecule total internal reflection fluorescence (smTIRF) microscopy, respectively, was unaffected over a wide range of ionic conditions. The extremely stable sliding clamp formed by MSH proteins on end-blocked DNA suggested that a fluorescence resonance energy transfer (FRET) system could be developed that was capable of clearly distinguishing HsMSH2-HsMSH3 and HsMSH2-HsMSH6 functions. We show that Cy5-labeled HsMSH2-HsMSH3 uniquely forms sliding clamps with a Cy3-labeled blocked-end (CA)_4_ lDL producing a strong time-averaged FRET signal. In contrast, Cy3-labeled HsMSH2-HsMHS6 appears more promiscuous but forms sliding clamps and time-averaged FRET more efficiently with Cy3-labeled DNA containing a G/T mismatch compared to the Cy3-labeled (CA)_4_ IDL. This FRET system is extremely durable and can tolerate room temperature manipulations and DMSO concentrations below 0.5%, a common solvent in high-throughput large-scale compound screens.

## RESULTS

### Ionic Dependence of Mismatch binding and ATP-induced Dissociation

MSH proteins preferentially bind DNA containing mismatched nucleotides (3, 15). Previous electrophoretic mobility shift analysis (EMSA) showed that binding to both a mismatched and fully duplex DNA increased with decreasing ionic strength (13,16,30). Similarly, kinetic analysis of MSH-mismatched binding has been examined by surface plasmon resonance (SPR), which is capable of distinguishing the second-order association rate (k_on_) and the first-order dissociation rate (k_off_) to generate equilibrium binding dissociation constants (*K_D_* = k_off_/k_on_) that are largely similar to EMSA (13,17,31).

SPR was utilized to determine that the *K_D_* for EcMutS (3.0 ± 0.4 nM and 1.5 ± 0.4 nM) and HsMSH2-HsMSH6 (0.7 ± 0.3 nM and 3.3 ± 2.3 nM) on unblocked versus biotin-streptavidin blocked-end mismatched DNA, respectively (**Supporting Fig. S1** and **S2**). These results confirm previous studies that demonstrated MSH binding to a mismatched was principally unaffected by whether the DNA end was open or blocked by a biotin-streptavidin linkage (13, 32).

We then examined the binding of EcMutS and HsMSH2-HsMSH6 to open and biotin-streptavidin blocked-end mismatched DNA by SPR over a range of ionic strength concentrations (**Fig. 1**; **Table 1**; **Supporting Table S2**). We observed very little variation in the *k_off_* for either open or blocked-end mismatched DNA (**Table 1**; **Supporting Table S2**). These results suggest that changes in mismatch binding (*K_D_*) by EcMutS and HsMSH2-HsMSH6 are mostly dependent on *k_on_* (*K_D_* ∼ C / *k_on_*, where C is a constant related to *k_off_*). Unlike previous electrophoretic mobility shift analysis (EMSA) in which MSH mismatch binding continually increased with decreasing ionic strength, we found that the *k_on_* for EcMutS and HsMSH2-HsMSH6 peaked at ∼75 mM NaCl (**Fig. 1**) (13,16,30). Calculating the apparent binding dissociation *K_D·app_* (*k_off_* / *k_on_*) did not change the shape of the curve since the *k_off_* increased by no more than by 20% at low NaCl concentrations, while the *k_on_* displayed more than a 2-fold change over the range of salt concentrations. These results appear to suggest that ionic strength influences MSH-mismatch binding differently when examined by the EMSA gel-based system compared to the SPR system that directly detects binding as surface mass changes.

**Table 1.** Dissociation Kinetics of EcMutS. Surface plasmon resonance dissociation studies. First-order rate constants of EcMutS (40 nM) in the absence (k_off_) and presence of ATP (1 mM, k_off ATP_). Open-end DNA (left) and blocked-end DNA (right) show similar rates of dissociation in the absence of ATP (k_off_). EcMutS sliding clamps quickly slide off open-end DNA (left), but are retained for (∼ 3 minutes) on blocked-end DNAs (right).

| **NaCl**<br>**(mM)** | **Open-End DNA** |  | **Blocked-End DNA** |  |
| --- | --- | --- | --- | --- |
| | $k_{\text{off}}^a$<br>($s^{-1}$) $\times 10^{-2}$ | $k_{\text{off} \cdot \text{ATP}}^b$<br>($s^{-1}$) $\times 10^{-2}$ | $k_{\text{off}}$<br>($s^{-1}$) $\times 10^{-2}$ | $k_{\text{off} \cdot \text{ATP}}$<br>($s^{-1}$) $\times 10^{-2}$ |
| 15 | 5.7 ± 0.19 | 5228 ± 943 | 5.9 ± 0.24 | 0.53 ± 0.01 |
| 30 | 5.9 ± 0.08 | 5028 ± 889 | 5.8 ± 0.10 | 0.58 ± 0.01 |
| 60 | 4.8 ± 0.17 | 5263 ± 233 | 5.4 ± 0.24 | 0.59 ± 0.03 |
| 80 | 5.1 ± 0.42 | 3931 ± 179 | 5.5 ± 0.14 | 0.67 ± 0.06 |
| 100 | 4.8 ± 0.12 | 2784 ± 51 | 5.3 ± 0.08 | 0.66 ± 0.04 |
| 120 | 5.0 ± 0.12 | 1557 ± 28 | 5.7 ± 0.10 | 0.86 ± 0.03 |
| 140 | 5.1 ± 0.28 | 1024 ± 147 | 5.5 ± 0.13 | 0.85 ± 0.04 |
| 160 | 5.4 ± 0.16 | 853 ± 64 | 6.2 ± 0.44 | 0.78 ± 0.01 |
| 180 | 5.4 ± 0.25 | 481 ± 69 | 6.6 ± 0.26 | 0.92 ± 0.04 |

<sup>a</sup> First-order rate constants in the absence of ATP (1 mM).
± Standard deviation of at least three replicates.
<sup>b</sup> First-order rate constants in the presence of ATP (1 mM).

**Figure 1.**
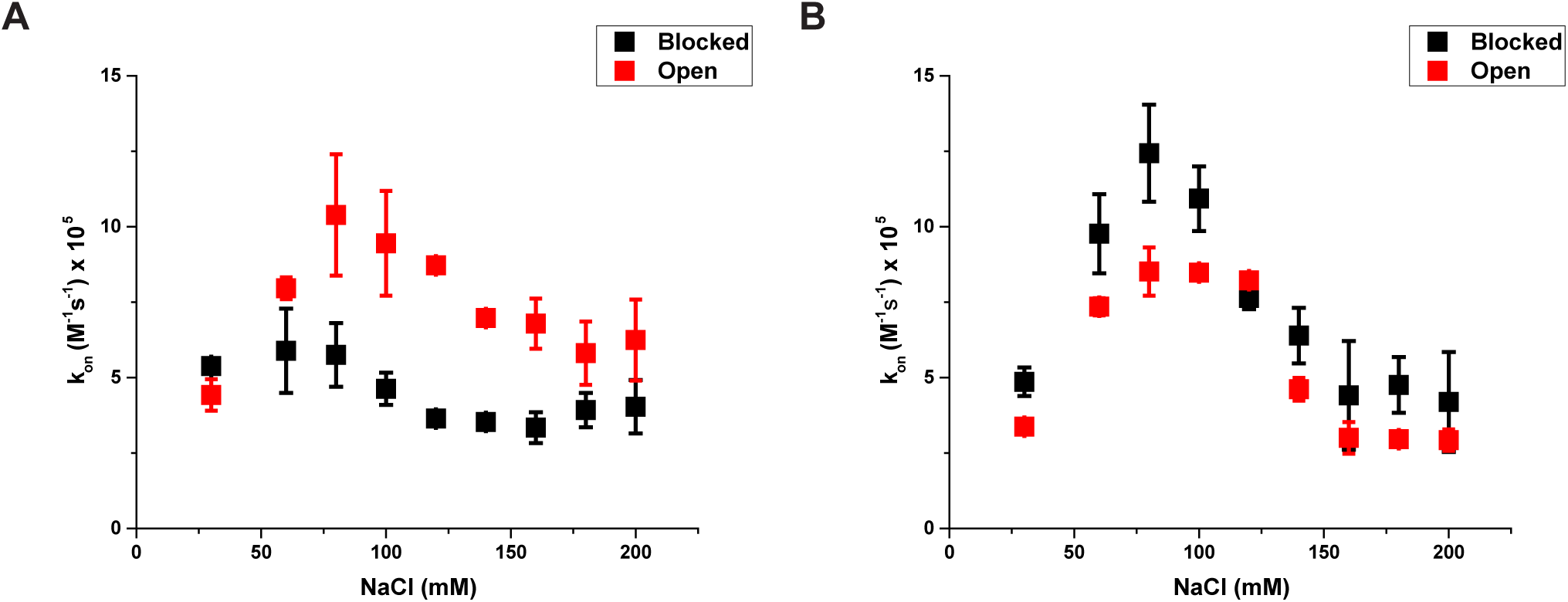
Binding Affinity of EcMutS and HsMSH2-HsMSH6. Surface plasmon resonance association rates. Second-order rate constants (*k_on_*) of (**A**) EcMutS (40 nM) and (**B**) HsMSH2-HsMSH6 (40 nM) in the absence of ATP across a wide range of salt concentrations. Blocked-end DNA is represented by black squares. Open-end DNA is represented by red squares. EcMutS and HsMSH2-HsMSH6 exhibit a peak binding affinity at ∼ 75 mM NaCl. Both EcMutS and HsMSH2-HsMSH6 *k_on_* curves display the same shape, but EcMutS has a higher binding affinity toward open-end DNA, likely due to an end binding effect.

We noted significant differences in the *k_on_* curves between open- and blocked-end mismatched DNA for both EcMutS and HsMSH2-HsMSH6, although the shapes appeared similar (**Fig. 1**). The largest difference occurred with EcMutS where the *k_on_* decreased as much as ∼2-fold with blocked-end mismatched DNA. These results appear to support anecdotal evidence that MutS homologs bind to oligonucleotide DNA ends when the mismatched DNA contains an open-end; a configuration that mimics double-stranded breaks (DSBs) which is rare *in vivo* (1).

The addition of ATP to an MSH protein bound to a mismatch or lesion provokes the formation of a sliding clamp that dissociates from an open-ended DNA (8,10,15,32). We examined the dissociation rate of EcMutS and HsMSH2-HsMSH6 bound to open- and blocked-end mismatched DNA by SPR following the addition of ATP (*k_off•ATP_*; **Table 1**; **Supporting Table S2**). As expected, the ATP-induced dissociation of EcMutS and HsMSH2-HsMSH6 from open-ended mismatched DNA was at least 10-fold faster than blocked-end mismatched DNA over a wide range of ionic strength conditions (**Table 1**; **Supporting Table S2**). These results are consistent with previous conclusions that ATP-bound MSH proteins rapidly dissociates from open DNA ends but are retained by blocked-end DNA (13,14,32). We note that the dissociation of EcMutS and HsMSH2-HsMSH6 in the absence of ATP on both open- and blocked-end mismatched DNA (*k_off_*) appeared similar to the *k_off•ATP_* with blocked-end mismatched DNA. While this rate equivalence may be a coincidence, it is possible that the dissociation mechanics of EcMutS and HsMSH2-HsMSH6 bound to a mismatch is similar to the dissociation mechanics of a trapped ATP-bound sliding clamp.

### ATP-bound MSH sliding clamps are extremely stable on mismatched DNA

Single molecule total internal reflection fluorescence microscopy (smTIRF) was used to probe the stability of Alexa647-labeled EcMutS sliding clamps on mismatched DNA (**Fig. 2**). In this system the trajectory, diffusion dynamics and lifetime of single EcMutS sliding clamps were determined over a range of ionic strengths. Numerous single particles could be easily resolved on the mismatched DNA (**Fig. 2A**; **Supporting Fig. S3**) and the trajectories binned to determine the diffusion coefficient as well as the lifetime. We found that the diffusion coefficient increased with increasing ionic strength (**Fig. 2B**). These observations are consistent with previous studies of *Thermus aquaticus* (Taq) TaqMutS and suggest that EcMutS ATP-bound sliding clamps undergo thermal diffusion along the DNA while maintaining intermittent contact with the backbone (14). In contrast, the lifetime appeared relatively constant (195 ± 11 sec) suggesting that EcMutS sliding clamps remain stably linked to the mismatched DNA over a wide range of ionic strength (**Fig. 2C**; **Supporting Fig. S4**).

**Figure 2.**
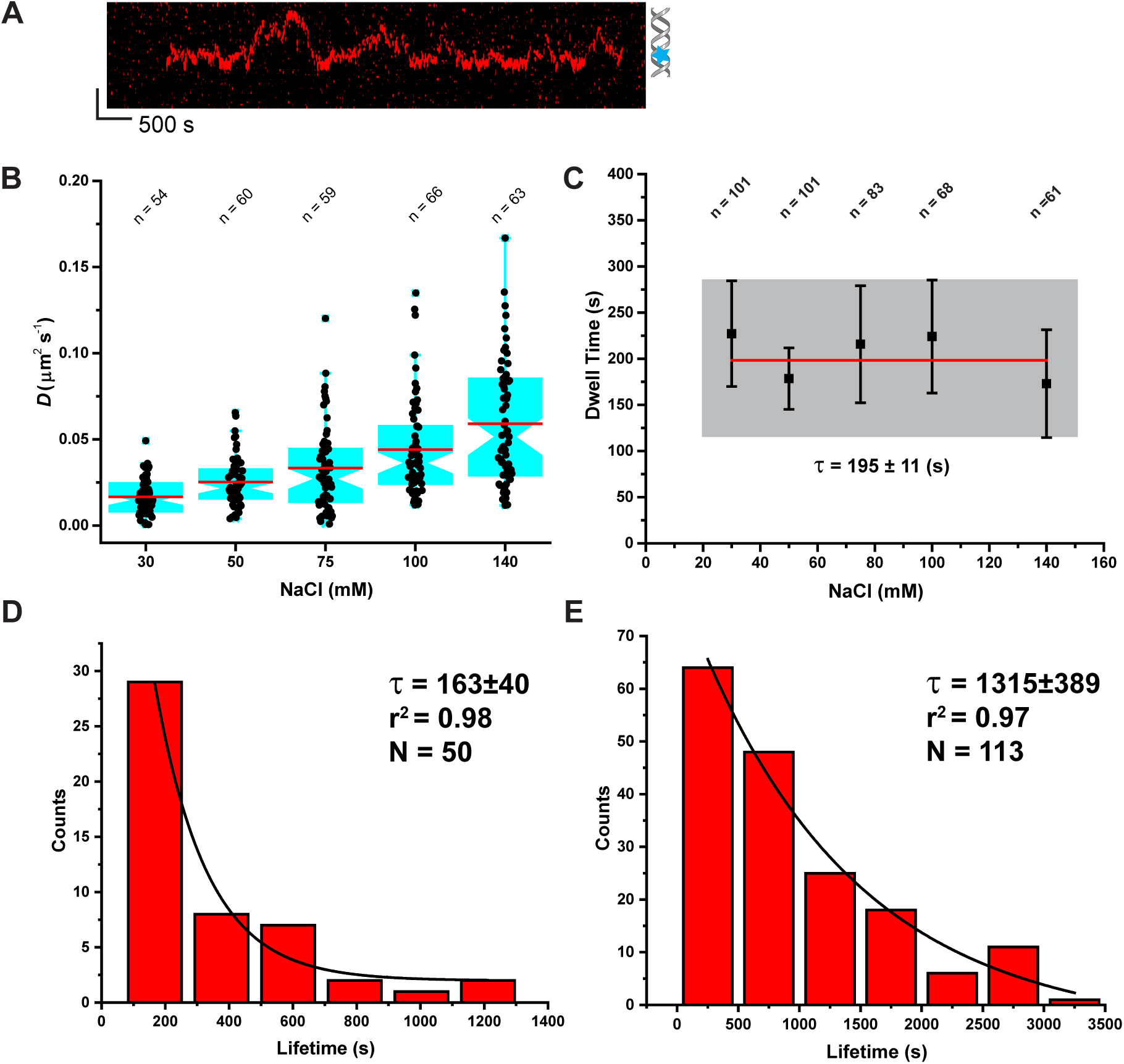
MSH Lifetimes. Single molecule analysis of individual labeled MSH sliding clamps. (**A**) Representative kymograph of EcMutS lifetime. Blue star indicates the mismatch position. (**B**) Box plots of diffusion for EcMutS (5 nM) sliding clamps in the prescence of ATP (1 mM) across a wide range of ionic strengths. Red line represents the mean, indent represents median. EcMutS sliding clamp diffusion rate increases with increasing NaCl concentration suggesting intermittent contact with the DNA backbone. (**C**)The lifetime of EcMutS (5 nM) sliding clamps in the presence of ATP (1 mM) across a wide range of ionic strength conditions. Black square represents average value, whiskers represent standard deviation. Average lifetime was found to be 195 ± 11 seconds. (**D**) The lifetime of HsMSH2-HsMSH6 (1 nM) sliding clamps in the presence of ATP (1 mM) at 100 mM NaCl. Average lifetime was found to be 163 ± 40 seconds. (**E**) The lifetime of HsMSH2-HsMSH3 (0.5 nM) sliding clamps in the presence of ATP (1 mM) at 100 mM KGlu. Average lifetime was found to be 1315 ± 389 seconds.

We similarly examine the lifetime of Cy3-labeled HsMSH2-HsMSH3 and HsMSH2-HsMSH6 sliding clamps on DNA by smTIRF (**Fig. 2D and 2E**, **Methods**). For these studies we used a G/T mismatch as the preferred substrate for HsMSH2-HsMSH6 and a CAG insertion-deletion loop-type (IDL) mismatch as the preferred substrate for HsMSH2-HsMSH3. While the lifetime of HsMSH2-HsMSH6 sliding clamps on DNA was similar to EcMutS (163 ± 40 sec; **Fig. 2D**), the lifetime of HsMSH2-HsMSH3 was almost six-times longer (1315 ± 389 sec; **Fig. 2E**). The analysis of HsMSH2-HsMSH3 was performed at different laser powers to assure photobleaching was not artificially reducing the lifetime (**Fig. 2E**, see colored histogram). Other than the IDL substrate preferences identified more than two decades ago (10), this is the first indication that HsMSH2-HsMSH3 may behave significantly different as a sliding clamp on the DNA compared to the major MSH proteins that are principally involved in post-replication MMR.

### A FRET-based system distinguishes human MSH sliding clamps

The stability of MSH sliding clamps on a mismatched DNA suggested that a bulk fluorescence resonance energy transfer (FRET) scheme might rapidly distinguish mismatch bound and sliding clamp forms of the human MSH heterodimer proteins. This system is based on previous smTIRF studies that determined the binding and sliding clamp kinetics of the *Thermus Aquaticus* MutS (TaqMutS) (12). We designed a 41-mer oligonucleotide containing either no mismatch, a G/T mismatch or a (CA)_4_ IDL with a biotin on both ends (**Fig. 3A**). These oligonucleotides were additionally labeled with Cy5 at a Thymine nucleotide nine base pairs from the mismatch (**Fig. 3A**). Previous studies with TaqMutS suggested that mispair-specific binding will place the Cy3-MSH in close proximity to the Cy5-DNA resulting in relatively high FRET. Because the Cy3-label is located on the N-terminus of HsMSH2 in the HsMSH2-HsMSH6 heterodimer and the N-terminus of HsMSH3 in the HsMSH2-HsMSH3 heterodimer, we expect a lower FRET efficiency than with the TaqMutS where the label was located on the clasp arm significantly closer to the Cy5-DNA label (12). In the presence of ATP, both HsMSH2-HsMSH3 and HsMSH2-HsMSH6 will form a sliding clamp (32, 33). However, when the glycosylation-free avidin derivative neutravidin is bound to the biotin ends of the DNA the ATP-bound MSH sliding clamps are projected to be retained where they will oscillate along the length of the oligonucleotide producing a time-averaged intermediate FRET (**Fig. 3A**).

**Figure 3.**
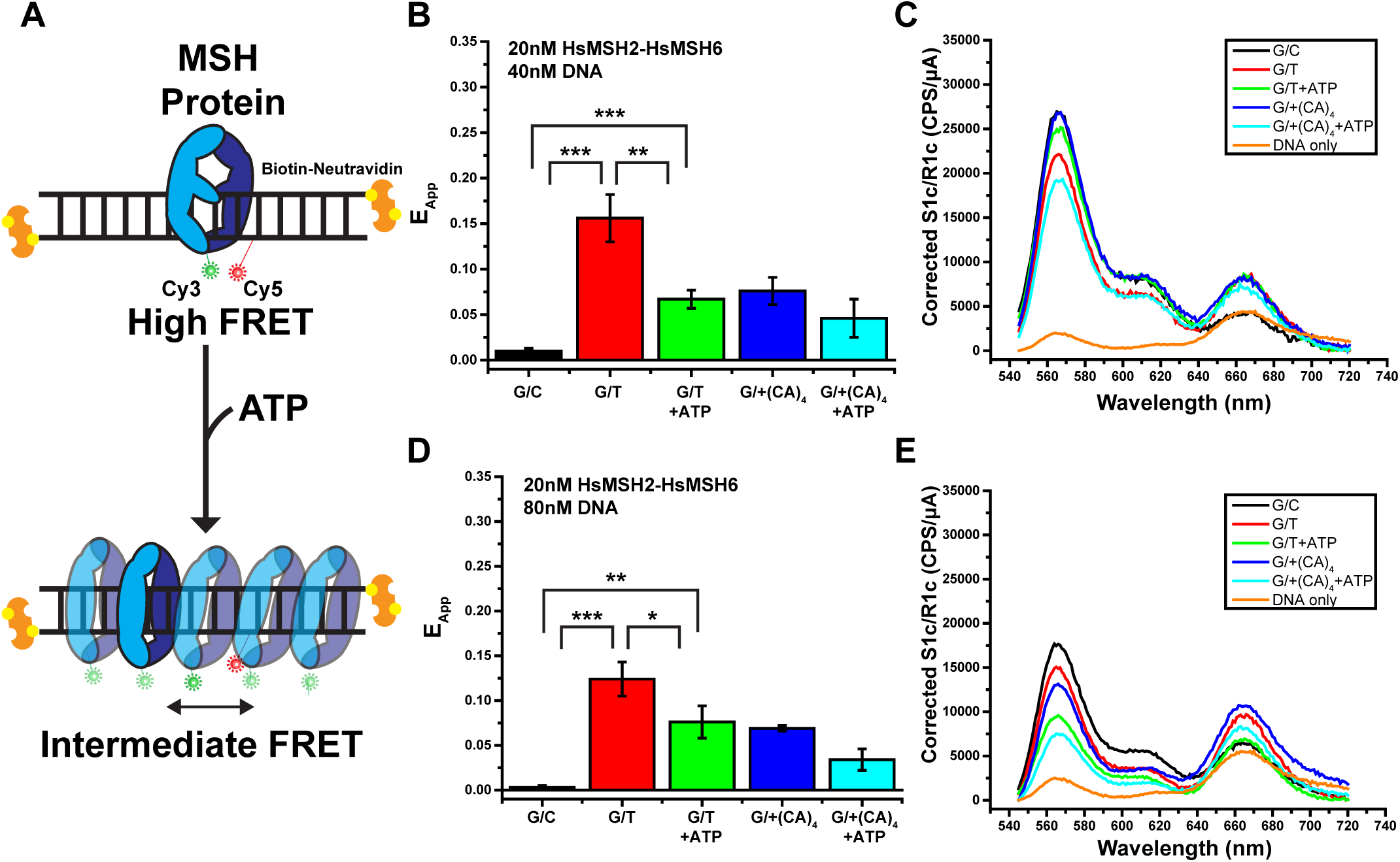
A FRET-Based Assay for Distinguishing HsMSH2-HsMSH6 and HsMSH2-HsMSH3 Activities. (**A**)The assay utilizes a short DNA oligo with double blocked-ends. These biotin-neutravidin linkages will trap MSH clamps on the DNA and prevent dissociation from the end. A FRET acceptor (Cy5) has been placed on the DNA near the mismatch. A FRET donor (Cy3) has been placed on the protein of interest. MSH protein binding to a mismatch results in a high FRET state (top). Upon the addition of ATP the MSH clamps will transition into stable sliding clamps that diffuse along the DNA. This conformation of the protein results in an intermediate or time averaged FRET (bottom) (**B**) E_App_ bar graph of 1:4 (HsMSH2-HsMSH6:DNA) for G/C; G/T; G/T + ATP;G/+(CA)_4_; G/+(CA)_4_ + ATP (mean ± SD). HsMSH2-HsMSH6 does not bind G/C DNA (black) and preferentially binds G/T DNA (red). HsMSH2-HsMSH6 displays a reduced FRET efficiency upon binding ATP. **, p < 0.01; p , 0.001; by unpaired *t* test. (**C**) Representative intensity vs. wavelength of 1:4 (HsMSH2-HsMSH6:DNA) for G/C; G/T; G/T + ATP; G/+(CA)_4_; G/+(CA)_4_ + ATP. (**D**) E_App_ bar graph of 1:2 (HsMSH2-HsMSH6:DNA) for G/C; G/T; G/T + ATP; G/+(CA)_4_; G/+(CA)_4_ + ATP (mean ± SD). HsMSH2-HsMSH6 does not bind G/C DNA (black) and preferentially binds G/T DNA (red). HsMSH2-HsMSH6 displays a reduced FRET efficiency upon binding ATP. *, p < 0.05; **, p < 0.01; ***, p < 0.001; by unpaired *t* test. (**E**) Representative intensity vs. wavelength of 1:2 (HsMSH2-HsMSH6:DNA) for G/C; G/T; G/T + ATP; G/+(CA)_4_; G/+(CA)_4_ + ATP.

A fixed concentration of HsMSH2-HsMSH6 (20 nM) was examined in the presence of a 1:2 and 1:4 molar ratio of Cy5-labeled DNA (**Fig. 3B and 3C**). As a control, we found that HsMSH2-HsMSH6 only transiently interacts with the G/C duplex DNA, resulting in little or no FRET (**Fig. 3B-E**). In contrast, HsMSH2-HsMSH6 specifically binds to the oligonucleotide containing a G/T mismatch resulting in a statistically significant elevated FRET (**Fig. 3B-E**; G/C vs G/T, p_40nM_ =0.0006, p_80nM_ =0.0004). As predicted, the addition of ATP resulted in a FRET reduction at both DNA concentrations (**Fig. 3B-E**; G/T vs G/T_ATP_, p_40nM_ = 0.005, p_80nM_ = 0.03), which remained significantly above the G/C background control (**Fig. 3B-E**; G/C vs G/T_ATP_, p_40nM_= 0.0007, p_80nM_ = 0.002). We noted a slight reduction in the G/T-binding FRET efficiency of HsMSH2-HsMSH6 with increased DNA concentration that is likely due to an increased background of direct Cy5 excitation by the 532 nm laser.

The *Saccharomyces cerevisiae* (Sc) ScMSH2-ScMSH6 heterodimer is known to recognize small IDLs in addition to single nucleotide mismatches (34). We found that incubation of the 8 bp (CA)_4_ IDL with HsMSH2-HsMSH6 resulted in significant FRET above the G/C control (**Fig. 2B-E**). Moreover, the FRET efficiency decreased in the presence of ATP consistent with the formation of a sliding clamp (**Fig. 2B-E**). In all cases, the FRET efficiency was reduced compared to the G/T mismatch (**Fig. 2B-E**). The observations are consistent with the conclusion that Cy3-labeled HsMSH2-HsMSH6 binds and forms an ATP-bound sliding clamp on Cy5-labeled mismatched DNA generating a strong and reproducible FRET. However, the FRET distinction between G/T and (CA)_4_ mismatched DNA by HsMSH2-HsMSH6 is limited.

Conversely, Cy3-labeled HsMSH2-HsMSH3 appears largely incapable of recognizing or forming sliding clamp with the G/T mismatched DNA resulting in FRET above the control G/C duplex DNA (**Fig. 3A-D**). Yet, a strong FRET was observed in the presence of the (CA)_4_ IDL mismatched oligonucleotide that resolved to a lower FRET in the presence of ATP (**Fig. 3A-D**). Compared to HsMSH2-HsMSH6 the distinction between G/T and (CA)_4_ IDL is significant. Moreover, the room temperature stability of HsMSH2-HsMSH3, where the smTIRF studies were performed, suggest that this FRET-based system might be useful as an initial inhibitory-compound screening tool.

### DMSO affects the FRET efficiency but is correctable

Most chemical compound libraries are dissolved in dimethyl sulfoxide (DMSO), a polar aprotic solvent capable of solubilizing polar and nonpolar compounds that is also miscible in water (35, 36). We examine the effect of DMSO on the protein and DNA fluorophore environment (**Fig. 4**). An intrinsic fluorescence increase was found around the FRET emission wavelengths (650-750 nm; **Fig. 4A**). Moreover, 5% DMSO significantly reduced the direct excitation and FRET excitation peaks when included with Cy3-labeled HsMSH2-HsMSH6 and Cy5-labeled G/T mismatched DNA. However, the decrease in Cy3 spectra appeared greater than the effect on the Cy5 spectra, which could dramatically affect the interpretation of FRET efficiency (**Fig. 4B**). We also noted a modest effect of DMSO on the direct excitation of Cy3-labeled HsMSH2-HsMSH6 and HsMSH2-HsMSH3 (**Fig. 4C,D**). Nevertheless, the overall shape of the Cy3 emission curve appeared largely unchanged suggesting that FRET excitation of a Cy5 fluorophore is achievable.

**Figure 4.**
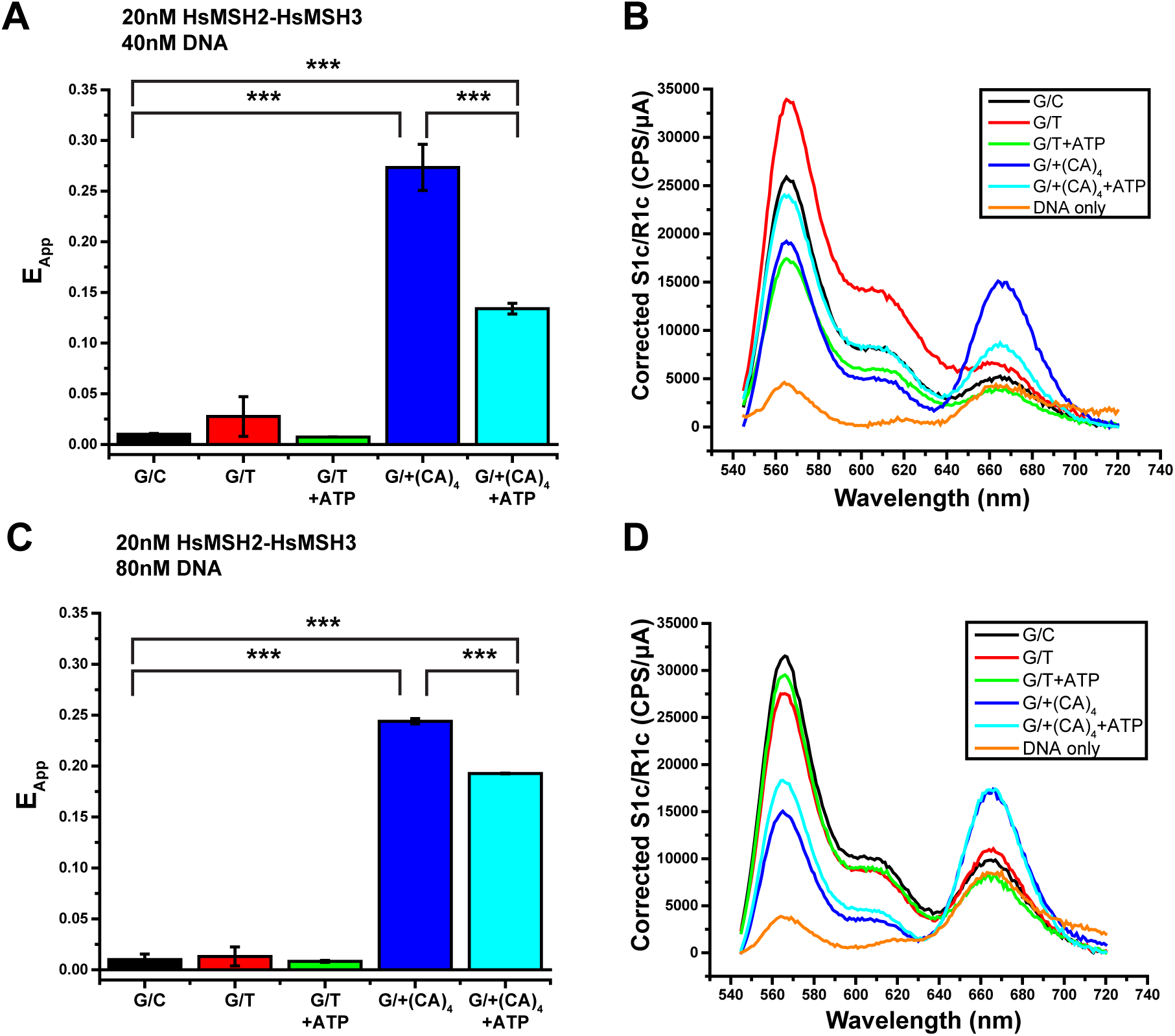
FRET Analysis of Mismatch Recognition by HsMSH2-HsMSH3. (**A**) E_App_ bar graph of 1:4 (HsMSH2-HsMSH3:DNA) for G/C; G/T; G/T + ATP; G/+(CA)_4_; G/+(CA)_4_ + ATP (mean ± SD). HsMSH2-HsMSH3 does not bind G/C DNA (black) and preferentially binds G/+(CA)_4_ DNA (red). HsMSH2-HsMSH3 displays a reduced FRET efficiency upon binding ATP. ***, p < 0.001; by unpaired *t* test. (**B**) Representative intensity vs. wavelength of 1:4 (HsMSH2-HsMSH3:DNA) for G/C; G/T; G/T + ATP; G/+(CA)_4_; G/+(CA)_4_ + ATP. (**C**) E_App_ bar graph of 1:2 (HsMSH2-HsMSH3:DNA) for G/C; G/T; G/T + ATP; G/+(CA)_4_; G/+(CA)_4_ + ATP (mean ± SD). HsMSH2-HsMSH3 does not bind G/C DNA (black) and preferentially binds G/(CA)_4_ DNA (red). HsMSH2-HsMSH3 displays a reduced FRET efficiency upon binding ATP. ***, p < 0.001; by unpaired *t* test(**D**) Representative intensity vs. wavelength of 1:2 (HsMSH2-HsMSH6:DNA) for G/C; G/T; G/T + ATP; G/+(CA)_4_; G/+(CA)_4_ + ATP.

The studies in the presence of 5% DMSO (**Fig. 4A,B**) implied that any interpretation of FRET efficiency must be performed in comparison to FRET values obtained in the presence of a G/C duplex DNA. Moreover, DMSO concentrations above 1% appeared to decrease the direct excitation of Cy3 significantly enough to reduce the FRET efficiency below detectable levels. Because HsMSH2-HsMSH6 displayed the lowest FRET efficiency compared to HsMSH2-HsMSH3, we used it as a bellwether to examine the effect of DMSO on the ability to distinguish mismatch binding (elevated FRET) and sliding clamp formation (intermediate-low FRET; **Fig. 5**). The G/T-binding FRET efficiency of HsMSH2-HsMSH6 decreased by at least 50% as the DMSO concentration was increased to 1% (**Fig. 5**). Nevertheless, these FRET efficiencies remained statistically significant compared to the G/C duplex DNA substrate (p_40nM_ = 0.0001, p_80nM_ = 0.01). Similarly, in the presence of ATP the FRET efficiency of sliding clamps bound to the blocked-end G/T mismatched DNA also decreased with increasing DMSO (**Fig. 5**). However, at both 40 nM and 80 nM DNA the FRET efficiency of a G/T mismatch in the presence of ATP was not significantly above the FRET efficiency of the G/C duplex DNA at concentrations of 0.5% DMSO and higher (**Fig. 5**). These observations are consistent with the conclusion that the MSH-DNA FRET-based system may be useful as an inhibitory-compound screening tool when the DMSO concentration is below 0.5%. Perhaps more importantly, it appears that this FRET system may be capable of differentially distinguishing the effects of inhibitory-compounds on HsMSH2-HsMSH6 and HsMSH2-HsMSH3 (discussed below).

**Figure 5.**
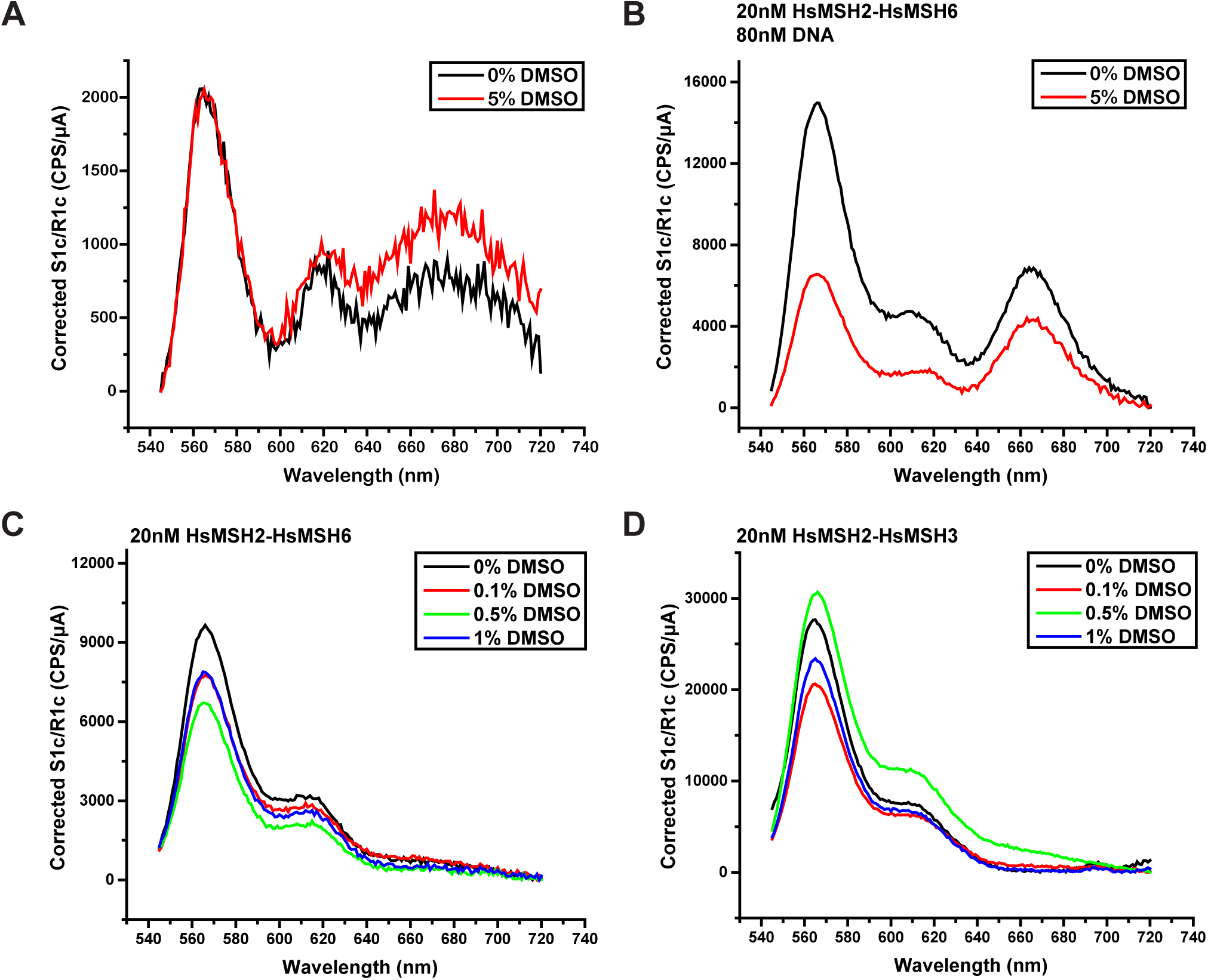
The Effect of DMSO on Fluorescence Excitation and Emission. Representative intensity vs. wavelengths of (**A**) DMSO in the absence of fluorophore. No effect is noted in the Cy3 spectra (550-650 nm). An increase in fluorescence is noted in the Cy5 spectra (650-750 nm). (**B**) DMSO in the presence of fluorophore. A decrease in the Cy3 spectra is observed, while an increase in the Cy5 spectra occurs. This decrease in donor (Cy3) and increase in acceptor (Cy5) will result in a skewed FRET value. (**C**) DMSO effect on HsMSH2-HsMSH6. (**D**) DMSO effect on HsMSH2-HsMSH3.

**Figure 6.**
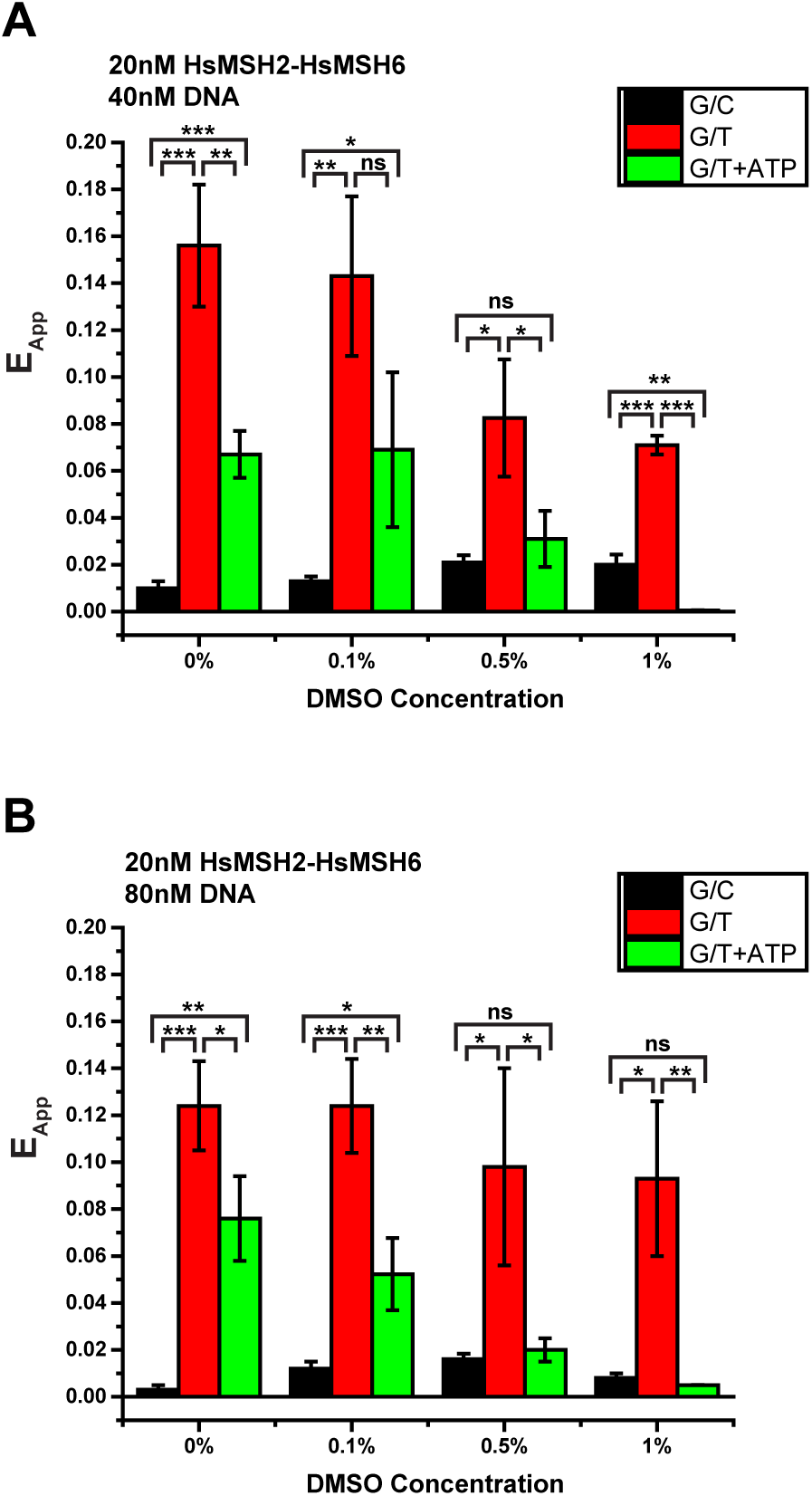
The Effect of DMSO on HsMSH2-HsMSH6 FRET. (**A**) E_App_ bar graph of 1:2 (HsMSH2-HsMSH6:DNA) for G/C; G/T; G/T + ATP in the presence of DMSO (mean ± SD). At 0.5% DMSO and greater the ability to distinguish between the 3 states (unbound (G/C), bound to a mismatch (G/T), sliding clamp (G/T + ATP)) is lost. *, p < 0.05; **, p < 0.01; ***, p < 0.001; by unpaired *t* test. (**B**) E_App_ bar graph of 1:4 (HsMSH2-HsMSH6:DNA) for G/C; G/T; G/T + ATP in the presence of DMSO (mean ± SD). At 0.5% DMSO and greater the ability to distinguish between the 3 states (unbound (G/C), bound to a mismatch (G/T), sliding clamp (G/T + ATP)) is lost. *, p < 0.05; **, p < 0.01; ***, p < 0.001; by unpaired *t* test.

## DISCUSSION

Decades of studies have clearly determined that MSH proteins recognized mismatched nucleotides and in the presence of ATP form an exceptionally stable sliding clamp on the DNA (8,10,12–14,16–18,32). This class of proteins includes members that appear principally involved in post-replication MMR as well as exclusive process such as those associated with double strand break (DSB) repair during meiosis (1, 37). In eukaryotes such as *S.cerevisae* and human cells, it is generally believed that two heterodimeric version, MSH2-MSH3 and MSH2-MSH6, play overlapping and redundant roles in post-replication MMR. However, it is worth noting that while MSH2-MSH6 appears quite promiscuous on recognizing all the single nucleotide mismatches as well as a variety of small-to-medium IDL mismatches (15, 38), MSH2-MSH3 only recognized single nucleotide mismatches poorly and clearly favors small-to-large IDL mismatches (10, 39). Moreover, the cellular concentration of MSH2-MSH3 appears to be at least 10-fold lower than MSH2-MSH6 (26, 40). Finally, no pathogenic mutations of *HsMSH3* have been found that lead to LS/HNPCC, while mutations of the core post-replication MMR genes *HsMSH2, HsMSH6, HsMLH1 and HsPMS2* have been identified as abundant causes of this cancer predisposition.

Using bulk biochemical and single molecule imaging analysis we have probed and compared the properties of *E.coli* EcMutS and human HsMSH2-HsMSH3 and HsMSH2-HsMSH6 proteins. All of these MSH proteins display exceptional stability as ATP-bound sliding clamps on DNA. Compared to TaqMutS (10 min), EcMutS (3.25 min) and HsMSH2-HsMSH6 (3.4 min) display a significantly shorter lifetime as sliding clamps on the mismatched DNA (**Fig. 2C,E**). However, it seems likely that the lifetime of TaqMutS might be inflated since ATP hydrolysis is required to release MSH sliding clamps from the DNA (12). Yet, Taq MutS is most active at temperatures above 40°C, while the single molecule imaging studies that are essential to generate lifetime data are performed at room temperature (∼23°C). Nevertheless, HsMSH2-HsMSH3 displayed a sliding clamp lifetime on the mismatched DNA of 17 min (**Fig. 2D**). This observation was a significant surprise and was confirmed at two different laser power settings to eliminate photobleaching, which could contribute to an inaccurate lifetime. Moreover, we note that the single molecule imaging studies supporting this long lifetime included time points in excess of 60 min, suggesting that HsMSH2-HsMSH3 remains active for an exceedingly long time at room temperature (∼23°C). Such protein stability is extremely important for inhibitory compound screening studies that utilize robotic distribution of components (see below). Taken as a whole, the extremely long lifetime of HsMSH2-HsMSH3 as a sliding clamp on DNA appears to widen the functional differences between it and the core MMR heterodimer HsMSH2-HsMSH6; potentially expanding and/or differentiating the cellular role(s) of HsMSH2-HsMSH3.

A major difference with MSH2-MSH6 is the role of MSH2-MSH3 in TNR expansion that leads to devastating diseases such as Huntington’s chorea and myotonic dystrophy (41). While mutation of the core MMR components that include MSH2-MSH6 results in expansions and contractions of short simple repeats (microsatellite instability) (42, 43), deletion of *MSH3* virtually eliminates TNR expansion (44, 45). These apparently opposing expansion-contraction phenotypes on repeat DNA sequences has never been adequately explained. One hypothesis for why *HsMSH3* mutations have not been linked to a cancer phenotype contemplates that a defect in HsMSH3 will lead to a significant increase in frame-shift mutations. These frameshifts would likely result in large numbers of abnormal or truncated surface antigens that could stimulate an immune response resulting in a termination of HsMSH3 mutant cells before a cancer phenotype could be established. Supporting this notion is the observation that PD1 inhibitors that reactivate immune recognition cells (46) are exceedingly effective therapeutic treatments for MMR defective tumors where frameshift mutations remain a minor but highly immunogenic component of the total mutation load (47).

These observations suggested that the identification of specific inhibitors of HsMSH2-HsMSH3 that have no effect on MSH2-MSH6 activities might be useful as therapeutics. Specific inhibition of HsMSH2-HsMSH3 could ameliorate the somatic effects that are generally the most immediately pathological in TNR diseases, while preventing the tumorigenesis associated with MMR defects. Our studies have reduced to practice a simple FRET-based screen that relies on the remarkably long lifetime and stability of MSH sliding clamps (**Fig. 3 and 4**). The assay relies on the conversion of mismatch binding events to a stable sliding clamp that continues to exhibit time-averaged FRET (**Fig. 3A**). We calculated the statistical effect size (z-factor) for high-throughput (HTP) drug screening comparing a variety of FRET-generating events (**Supporting Fig. S5**) (48, 49). What is most evident from these plots is that FRET events generated by HsMSH2-HsMSH3 provide the largest z-factor margin for the detection of inhibitory events. These calculations suggest that in initial HTP compound screen focused on HsMSH2-HsMSH3 functions should provide an array of specific inhibitors. This initial screen could then be followed by a counter screen for HsMSH2-HsMSH6 functions to identify specific inhibitors of HsMSH2-HsMSH3 that do not affect HsMSH2-HsMSH6. Because the z-factor for HsMSH2-HsMSH6 functions in significantly lower than HsMSH2-HsMSH3 functions, more redundant FRET repeats will be required to assure statistical significance (**Supporting Fig. S5**). Nevertheless, the stability of the FRET system should be able to easily accommodate the extra duplicates.

## EXPERIMENTAL PROCEDURES

### EcMutS expression, purification, and labelling

EcMutS protein was expressed, purified, and labeled with fluorophore as previously described (50). Briefly, EcMutS was expressed in BL21 AI E. coli cells and purified by FPLC using a Ni-NTA column (Qiagen) and Heparin (GE Healthcare) column (18). EcMutS-containing fractions were dialyzed in storage buffer (25 mM HEPES pH 7.8, 1 mM DTT, 0.1 mM EDTA, 150 mM NaCl, 20 % glycerol) and frozen at -80°C.

For fluorophore labeled proteins, EcMutS and MtFGE (ratio 1:1) were dialyzed together in conversion buffer at 4°C for 48 hr and then dialyzed in labeling buffer overnight. Proteins were then incubated with HIPS-AlexaFluor 647 at 0°C for 48 hr (50). Labelled EcMutS was separated from free dye and MtFGE on a heparin column. Fractions were visualized on an 8% SDS-PAGE and dialyzed in storage buffer (25 mM HEPES pH 7.8, 1 mM DTT, 0.1 mM EDTA, 150 mM NaCl, 20 % glycerol) and frozen at -80°C.

### HsMSH2-HsMSH6 expression and purification

HsMSH2-MSH6 protein was expressed and purified as previously described (51). Briefly, HsMSH2 and HsMSH6 were coexpressed in Hi5 insect cells using the Bac-to-Bac Baculovirus Expression System (Invitrogen). Protein was purified by FPLC using a Ni-NTA column (Qiagen), PBE-94 column (Sigma), HiLoad 16/60 Superdex 200 (GE Healthcare), and MonoQ 5/50 GL (GE Healthcare). HsMSH2-MSH6 containing fractions were dialyzed in storage buffer (25 mM HEPES pH 7.8, 1 mM DTT, 0.1 mM EDTA, 150 mM NaCl, 20 % glycerol) and frozen at -80°C.

### DNA preparation

For surface plasmon resonance DNAs (82-mers) were prepared as follows. Oligonucleotides were synthesized by Integrated DNA Technologies (Coralville, Iowa; Supplementary Table S1). The single stranded (ss) oligos were purified by polyacrylamide gel electrophoresis (PAGE), and annealed overnight by step-down cooling in a thermocyler to create double stranded (ds) oligos. The G-Strand used in every construct contained the biotin moiety that is necessary for surface linkage (**Supplementary Table S1**). The complementary T-strand either contained a 5’ digoxigenin (dig) modification or no modification (**Supplementary Table S1**) and when annealed to the G-strand created a G/T mismatch. The annealed oligos were purified using a GenPak Fax (Waters) column on an HPLC.

The DNA for smTIRF was prepared as previously described (18). Briefly, a 7-kb fragment of DNA was prepared by digesting a plasmid with *BsaI*. λ-phage DNA (3.2 nM, Thermo Scientific) was ligated with Lambda Mismatch 2A and Lambda Mismatch 2B (800 nM, **Supplementary Table S1**) followed by digestion with *BsaI*. The treated λ-DNA was then ligated with the 7-kb DNA, 1000x Lambda Hairpin Linker 1 and Lambda Hairpin Linker 2 (**Supplementary Table S1**). The 18.4-kb band was excised and purified utilizing β-agarase, ethanol precipitation and stored at -80°C (18).

The 41-mer oligonucleotides for FRET studies were synthesized by Integrated DNA Technologies (Coralville, Iowa) and double stranded oligos were prepared by annealing the G-Strand with either the T-Strand (G/T mismatch) or the C-Strand (homoduplex; **Supplementary Table S1**). All strands contained 3’ biotin moieties. The oligos were annealed overnight in a thermocycler by step-down cooling. The annealed oligos were purified from contaminates using a GenPak Fax column (Waters) by HPLC.

### Preparation of surface plasmon resonance (SPR) sensor chip

A streptavidin coated Biacore 3000 SPR sensor chip (Sensor Chip SA, GE Healthcare) containing four channels was preconditioned with NaOH (50 mM). Biotin only and biotin-digoxigenin dsDNA (described above) was immobilized on the chip, with dsG/T-dig (blocked-end) DNA in channel 2 and dsG/T (open-end) DNA in channel 4, leaving channels 1 and 3 as buffer-only reference channels. The SPR response units (RUs) were kept within 10 between channels 2 and 4 to ensure the similar amounts of DNA was immobilized in each channel.

### SPR analysis of MSH activity

The SPR flow scheme is outlined in **Supplementary Figures S1** and **S2**. Briefly, the MSH protein was injected to monitor binding (k_on_) followed by a buffer wash step that monitored dissociation (k_off_). The binding-dissociation was followed by injection of binding-dissociation buffer containing ATP that results in a transition to an MSH sliding clamp that rapidly dissociated from the open end of an unblocked mismatched DNA (k_off•ATP_; **Supplementary Figures S1B** and **S2B**). In contrast, blocked-end mismatched DNA retains the MSH sliding clamps that exhibit a dissociation rate which reflects intrinsic stability (k _off•SC_; **Supplementary Figures S1A** and **S2A**). Following the ATP injection, the surface was regenerated with 1 M NaCl to remove any remaining protein while leaving the DNA undamaged.

For ionic strength binding/stability analysis, a titration of EcMutS and HsMSH2-MSH6 (0 – 100 nM) was first performed to determine the protein concentration that reflected ∼80% binding saturation. The experimental concentration for both MSH proteins (40 nM) for ionic strength binding/stability analysis fit to a Langmuir curve. Injections containing no protein were performed at the beginning and end of the analysis to ensure that background binding was unchanged between analysis.

### SPR analysis

Experiments were performed at 25°C. EcMutS and HsMSH2-MSH6 were examined independently using the same chip. A pre-blocking of the dig with anti-digoxigenin (anti-dig, 50 nM, Roche) was performed prior to each experiment. While no change in SPR RUs was noted, previous work has shown that the anti-digoxigenin is enough to block/trap MSH clamps on the DNA (51, 52). The running buffer for both proteins was 25 mM HEPES pH 7.8, 0.1 mM DTT, 0.1 mM EDTA, 10 mM MgCl_2_, 200 µg/ml acetylated BSA (Promega), 0.01% P-20 surfactant (GE Healthcare), 1 nM anti-dig (Roche), and stated [NaCl]. Proteins were diluted prior to experiment to reduce glycerol and adjust NaCl concentrations. Binding experiments consisted of ten NaCl conditions 15, 30, 60, 80, 100, 120, 140, 160, 180, 200 mM NaCl. When ionic strength is referred to it is only that of NaCl. RU changes reflect the binding and dissociation over time. ATP-induced dissociation was tested with an addition of running buffer containing ATP (1 mM, Roche) following binding of the MSH homolog.

Raw RUs were collected for each independent channel (1–4). Subtractions of the empty reference channels (2-1, 4-3) was performed to account of non-specific adsorption of protein to the surface. This subtraction data was used for all analysis. Data was normalized to the maximum of exponential fits for MSH binding. Runs were performed in triplicate. Association and dissociation constants were determined by fitting an exponential decay equation (Origin Labs) to the appropriate section of the curve.

### Preparation of smTIRF flow cell

A laboratory engineered flow cell was used for all experiments consisting of a neutravidin coated, PEG passivated quartz slide surface. Laminar flow (250 μL/min) was used to inject and stretch the biotinylated dsDNA (as described above, 300 pM) in 300 μL T-50 buffer (20 mM Tris-HCL, pH 7.5, 50 mM NaCl) on the surface. Prior to performing an experiment, the flow cells and all relevant tubing were prepared with BSA and P-20 to prevent nonspecific adsorption that may reduce concentrations.

### smTIRF experiments and data analysis

Single molecule fluorescence data was attained on a home-built prism-type TIRF microscope (18) with green (532 nm) and red (635 nm) laser lines. Emissions were split by a Dual View optical setup (DV2, Photometrics) in bypass mode before collection using an EMCCD camera (ProEM Exelon512, Princeton Instruments). Laser excitation was modulated by the opening and closing of shutter controlled by Micro-Manager image capture software (53).

The single molecule imaging buffer was 20 mM Tris-HCl (pH 7.5), 5 mM MgCl_2_, 0.1 mM DTT, 200 µg/ml acetylated BSA (Promega), 0.0025% P-20 surfactant (GE Healthcare), 1 mM ATP (Roche), and stated [NaCl]. The imaging buffer included saturated (∼3 mM) Trolox, PCA (1 mM), and PCD (10 nM) to minimize photo blinking and photobleaching (54, 55).

EcMutS lifetime was calculated by flowing AF647-EcMutS (5 nM) into the prepared flow cell. In the absence of flow protein-DNA interactions were observed live. After recording Syto 59 (1000 nM, Invitrogen) was used to stain the DNA.

### Preparation of the Cy3-peptide

10 mg of lyophilized peptide containing a Sortase recognition sequence(56) (57) and a single cystine residue for maleimide labeling (CLPETGG, GenScript) was dissolved in reaction buffer (50 mM Tris 7.0 and 5 mM TCEP) and incubated at room temperature for 30 min. The reaction was then added to 3 mg of lyophilized Sulfo-Cy3 maleimide (Lumiprobe) and incubated overnight at 4°C. The labeled peptide was purified with reverse phase HPLC chromatography (Zorbax SB-C18, Agilent). Peak fractions were lyophilized and stored at -80°C until needed. For labeling the peptide was dissolved in Sortase reaction buffer (25 mM HEPES pH 7.8, 150 mM NaCl and 10% glycerol).

### MSH Sortase-tag addition, expression, purification, and labeling

MSH proteins may be modified on the N-terminus to contain a hexa-histidine (his_6_), two serine spacers, a Sortase recognition sequence, and a flexible linker (GGGS) attached to the MSH protein of interest. For these studies the tags were placed on HsMSH6 and HsMSH3. MSH proteins were cloned into pFastBac1 (Invitrogen). The heterodimeric proteins (HsMSH2-HsMSH6, HsMSH2-HsMSH3) were coexpressed as previously described (10,25,32) in Sf9 cells using the Bac-to-Bac Baculovirus Expression System (Invitrogen). Protein was first purified by FPLC using a Ni-NTA column (Qiagen) followed by a heparin column (GE Healthcare). Peak fractions were pooled and combined with 4-times molar ratio of the Sortase protein and 50-times molar ratio of the Cy3-Peptide for 30 min at 4°C (56, 58). The reaction is quenched with 20 mM EDTA and loaded onto a spin-desalting column (40K Zeba Spin Desalting Column, Thermo Scientific). The eluent was then separated from the Cy3-peptide using a second heparin column (GE Healthcare) and polished using a MonoQ 5/50 GL (GE Healthcare). The HsMSH2-HsMSH6 and HsMSH2-HsMSH3 containing fractions were dialyzed in storage buffer (25 mM HEPES pH 7.8, 1 mM DTT, 0.1 mM EDTA, 150 mM NaCl, 20 % glycerol) and frozen at -80°C.

### DNA fluorophore labeling and preparation

DNA oligonucleotides are labeled with near 100% efficiency using fluorescent (Cy/AF) dyes. To increase homogeneity between multiple substrates, the G-strand is always labeled with the acceptor fluorophore and then annealed with a complementary strand containing no mismatch (G/C), a single nucleotide mismatch (G/T) or an IDL mismatch [G/+(CA)_4_].

Single strand DNA containing an amino modifier C6 dT (IDT) in aqueous solution (10mM Tris-HCl pH 8.0, 1mM EDTA) was precipitated using 3M sodium acetate pH 5.2, 95% ethanol, and glycogen, followed by a wash with 70% ethanol to remove any impurities remaining from the synthesis process (**Supplementary Table S1**). The precipitate was then air-dried and resuspended in labeling buffer (70-100ul 0.1M sodium tetraborate pH 8.5). Cy-dye or analog, dissolved in dimethylformamide and added to the reaction in 10-30x molar excess. The reaction was mixed until completely dissolved and kept in foil rotating overnight at 23^°^C and 500 rpm. The overnight labeled material was ethanol precipitated following the same protocol as above twice to remove free dye. The final pellet should be fluorescent and if color is not observed the labeling should be repeated before subsequent HPLC purification steps. The final pellet was dissolved in triethylamine acetate (TEAA).

The fluorophore-labeled single-stranded DNA oligonucleotide was separated from unlabeled oligonucleotide and unincorporated free fluorophore by C18 reverse-phase HPLC chromatography (Poroshell 120 EC-C18, Agilent). Labeled fractions were pooled and ethanol precipitated as described above and resuspended in aqueous solution (10mM Tris-HCl pH 8.0, 1mM EDTA).

The fluorophore-labeled G-strand oligonucleotide was annealed with complementary unlabeled DNA (C-strand no mismatch, T-strand single nucleotide mismatch, +(CA)_4_-strand IDL mismatch) in a 1:1 molar ratio by heating to 95°C and slow step cooling (**Supplementary Table S1**). The duplex DNA products were then purified from any remaining single stranded DNA substrates by ion exchange HPLC chromatography (Gen-Pack Fax, Waters). Duplex fractions were pooled, and ethanol precipitated as described above and resuspended in aqueous solution (10mM Tris-HCl pH 8.0, 1mM EDTA).

### FRET mismatch recognition assay analysis

FRET detection was performed utilizing a FlouroMax-4 (Horiba Jobin Yvon) according to manufacturer’s recommendations (59). The Flouromax-4 (Horiba Jobin Yvon) is a corrected photon counting system where the Intensity (counts per sec / μA) is: corrected signal detector (S1c) / corrected reference detector (R1c) or S1c/R1c. Because FRET calculations are ratios of acceptor and donor intensities, the S1c/R1c read-out from the FluoroMax (or any other corrected photon counting reader) may be used directly as measures of fluorescence intensity (I). While peak intensity at specific donor (∼560 nm) and acceptor (∼670 nm) wavelengths may be used as an initial screen for FRET efficiency calculations, increased accuracy is obtained by fitting the entire scanned peak intensities to a Gaussian function and integrating. Apparent FRET efficiency (E_App_) is calculated by: E_App_ = (I_A_ - I_A•DNA Only_) / [(I_A_ - I_A•DNA Only_) + (I_D_ - I_D•DNA Only_)]; where I_A_ is the MSH+DNA acceptor intensity, I_A•DNA Only_ is the acceptor intensity of the DNA alone, I_D_ is the donor intensity, and I_D•DNA Only_ is the donor intensity of the DNA alone. Thus, E_App_ corrects for background contribution of 510 nm excitation that results in the acceptor Cy5-DNA Gaussian emissions with peaks at ∼560 nm and at ∼670 nm in the absence of donor Cy3-MSH protein. The donor Cy3-MSH protein alone does not contribute to background Gaussian emission with a peak at ∼670 nm, and therefore does not require a correction factor.

The emission spectra was scanned from 545 nm to 720 nm following 510 nm excitation with a xenon lamp. The Cy3 peak was fit with 2 Gaussian functions due to the emission spectra of Cy3. The Cy5 emission peak was fit with a single Gaussian function. The E_App_ FRET was calculated by measuring energy transfer under donor/acceptor intensity ratio (ratiometric FRET). A Cy5 labeled 41-bp duplex DNA containing a G/C duplex, G/T mismatch, or +(CA)_4_ IDL mismatch with 3’ biotin on both strands was end blocked with neutravidin was used. Cy3 labeled MSH protein was kept constant at 20 nM. The molar ratio was varied by increasing and decreasing the amount of DNA. Experiments were performed in a quartz cuvette on a FlouroMax-4 (Horiba Jobin Yvon) 5 minutes following the addition of protein. The addition of 1 mM ATP was tested. Binding of MSH protein to the mismatch should result in a high FRET value (**Figure 1**, left). The addition of ATP should result in a sliding clamp that results in a time-averaged FRET (**Figure 1**, right). As the ratio of donor to acceptor is increased a higher FRET value is observed.

## SUPPORTING INFORMATION

Supporting Information are available online.

## AUTHOR CONTRIBURIONS

B.M.B., J-B.L., and R.F. designed the experiments. B.M.B, J.L., J.M-L., and J.A.L. purified and labeled the proteins. B.M.B. purified the DNA substrates and performed surface plasmon resonance, single-molecule studies, and fluorimeter assays. B.M.B., N.D.J., J-B.L., and R.F. analyzed the data and wrote the paper. All authors participated in critical discussions.

## ACKNOWLEDGEMENTS

We would like to thank members of the Fishel laboratory for insights and helpful discussions.

## FUNDING

This work was supported by National Institutes of Health grants CA067007 and GM129764 (R.F.) and the Global Research Lab Program through the NRF of Korea funded by the Ministry of Science and ICT 2017K1A1A2013241 (J.-B.L.).

---

**Supporting Table S1.**
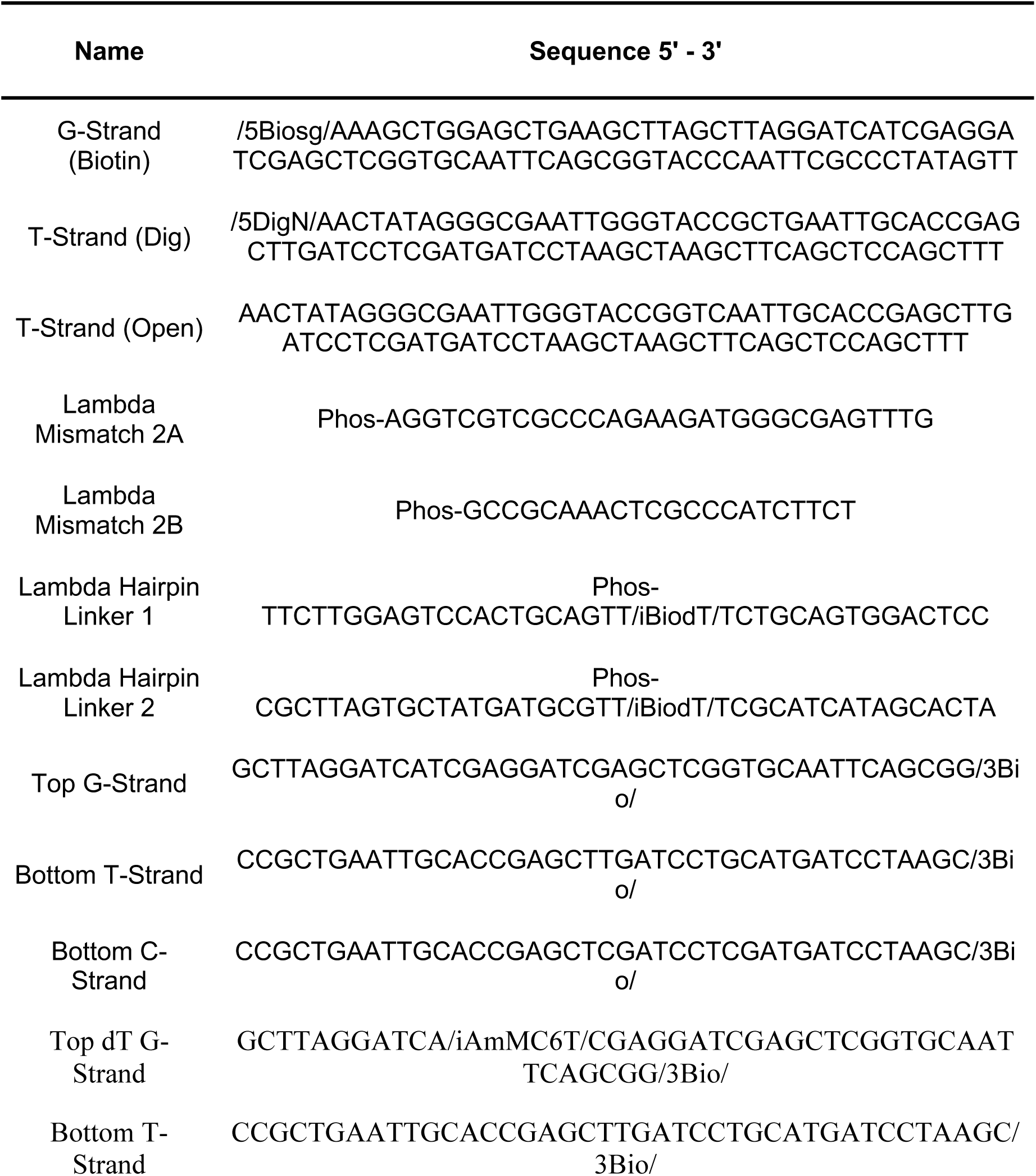
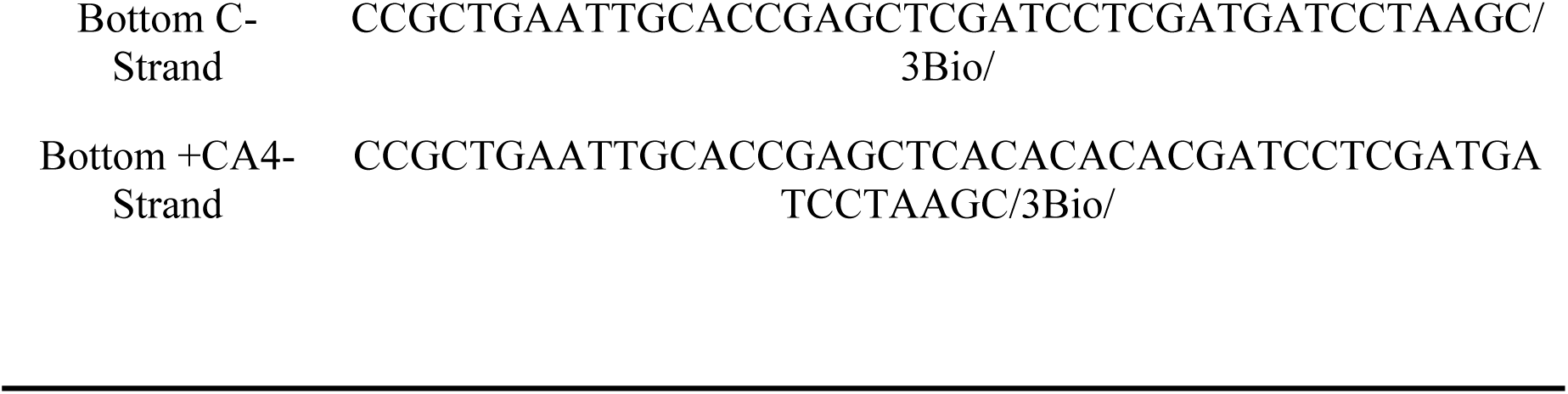

**Supporting Table 2.** Dissociation Kinetics of HsMSH2-HsMSH6. Surface plasmon resonance dissociation studies. First-order rate constants of HsMSH2-HsMSH6 (40 nM) in the absence (k_off_) and presence of ATP (1 mM, k_off ATP_). Open-end DNA (left) and blocked-end DNA (right) show similar rates of dissociation in the absence of ATP (k_off_). HsMSH2-HsMSH6 sliding clamps quickly slide off open-end DNA (left) but are retained for ∼ 3 minutes on blocked-end DNAs (right).

| NaCl<br>(mM) | Open-End DNA |  | Blocked-End DNA |  |
| --- | --- | --- | --- | --- |
| | $k_{\text{off}}^a$<br>($s^{-1}$) $\times 10^{-2}$ | $k_{\text{off}\cdot\text{ATP}}^b$<br>($s^{-1}$) $\times 10^{-2}$ | $k_{\text{off}}$<br>($s^{-1}$) $\times 10^{-2}$ | $k_{\text{off}\cdot\text{ATP}}$<br>($s^{-1}$) $\times 10^{-2}$ |
| 15 | 0.62 ± 0.30 | 20.3 ± 0.57 | 1.1 ± 0.44 | 0.47 ± 0.01 |
| 30 | 0.03 ± 0.01 | 24.5 ± 1.2 | 0.54 ± 0.07 | 0.54 ± 0.02 |
| 80 | 0.88 ± 0.12 | 26.7 ± 0.77 | 1.4 ± 0.15 | 0.35 ± 0.01 |
| 100 | 0.61 ± 0.02 | 26.6 ± 0.93 | 1.6 ± 0.09 | 0.56 ± 0.00 |
| 120 | 0.84 ± 0.07 | 28.8 ± 1.2 | 2.2 ± 0.13 | 0.55 ± 0.00 |
| 140 | 1.3 ± 0.09 | 30.3 ± 1.5 | 2.8 ± 0.80 | 0.52 ± 0.01 |
| 160 | 1.2 ± 0.12 | 23.6 ± 1.3 | 4.6 ± 0.88 | 0.52 ± 0.01 |
| 180 | 0.92 ± 0.05 | 19.0 ± 0.68 | 6.3 ± 1.3 | 0.60 ± 0.01 |

<sup>a</sup> First-order rate constants in the absence of ATP (1 mM).
± Standard deviation of at least three replicates.
<sup>b</sup> First-order rate constants in the presence of ATP (1 mM).

**Supporting Figure S1.**
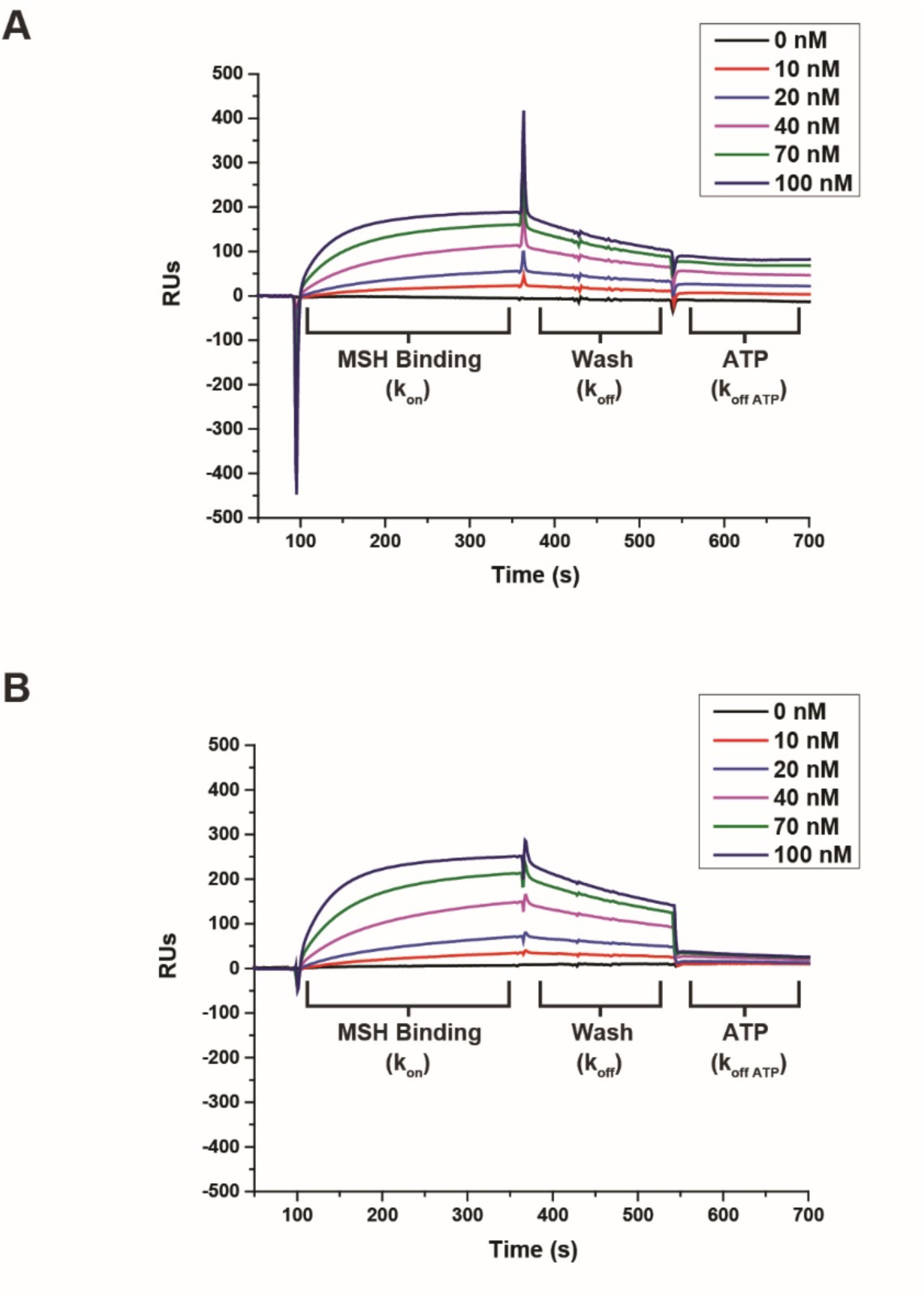
EcMutS Binding Analysis. Surface plasmon resonance association studies. (**A**) Protein titration of EcMutS on blocked-end DNA. (**B**) Protein titration of EcMutS on open-end DNA. MSH Binding indicates where MSH (40 nM) protein is flowed onto chip. This region is used to calculate the second-order rate constant (*k_on_*). Wash indicates where the unbound protein is washed from chip. This region is used to calculate the first-order rate constant (*k_off_*). ATP indicates where ATP (1 mM) in buffer is flowed onto chip. This region is used to calculate the first-order rate constant (*k_off·ATP_*).

**Supporting Figure S2.**
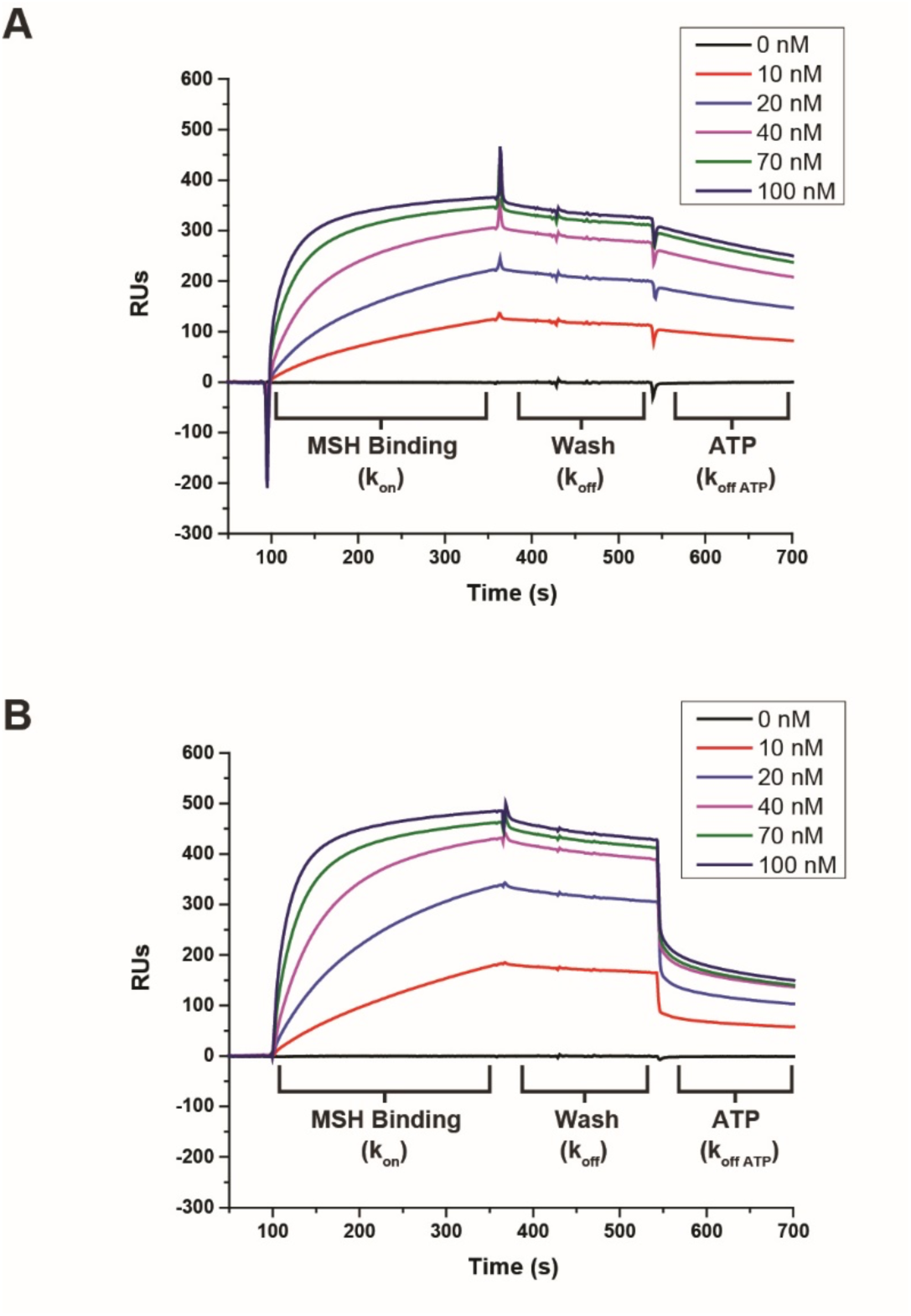
HsMSH2-MSH6 Binding Analysis. Surface plasmon resonance association studies. (**A**) Protein titration of HsMSH2-MSH6 on blocked-end DNA. (**B**) Protein titration of HsMSH2-MSH6 on open-end DNA. MSH Binding indicates where MSH (40 nM) protein is flowed onto chip. This region is used to calculate the second-order rate constant (*k_on_*). Wash indicates where the unbound protein is washed from chip. This region is used to calculate the first-order rate constant (*k_off_*). ATP indicates where ATP (1 mM) in buffer is flowed onto chip. This region is used to calculate the first-order rate constant (*k_off·ATP_*).

**Supporting Figure S3.**
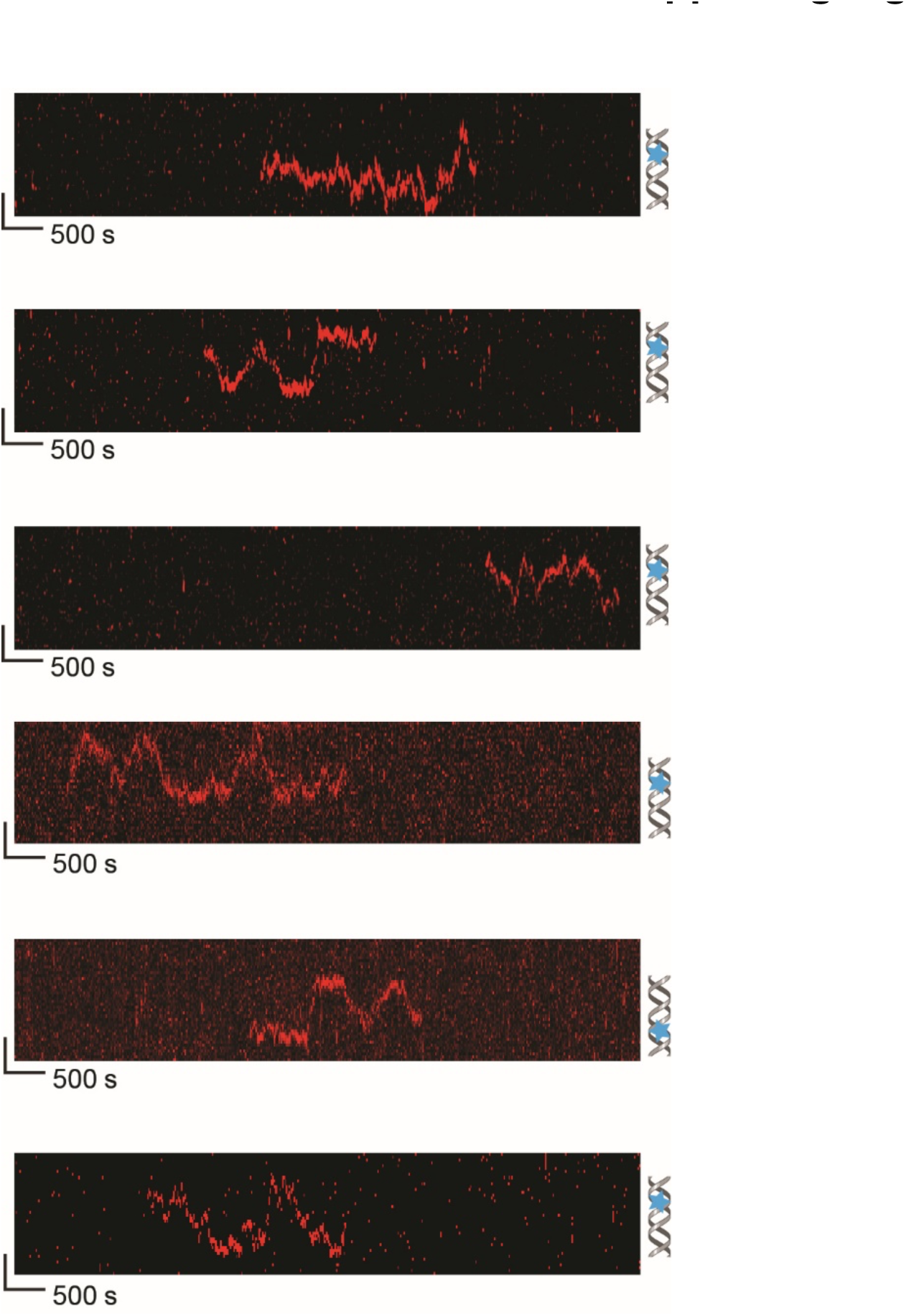
Additional MutS Kymographs. Representative kymographs of EcMutS DNA search. Blue star indicates the mismatch position.

**Supporting Figure S4.**
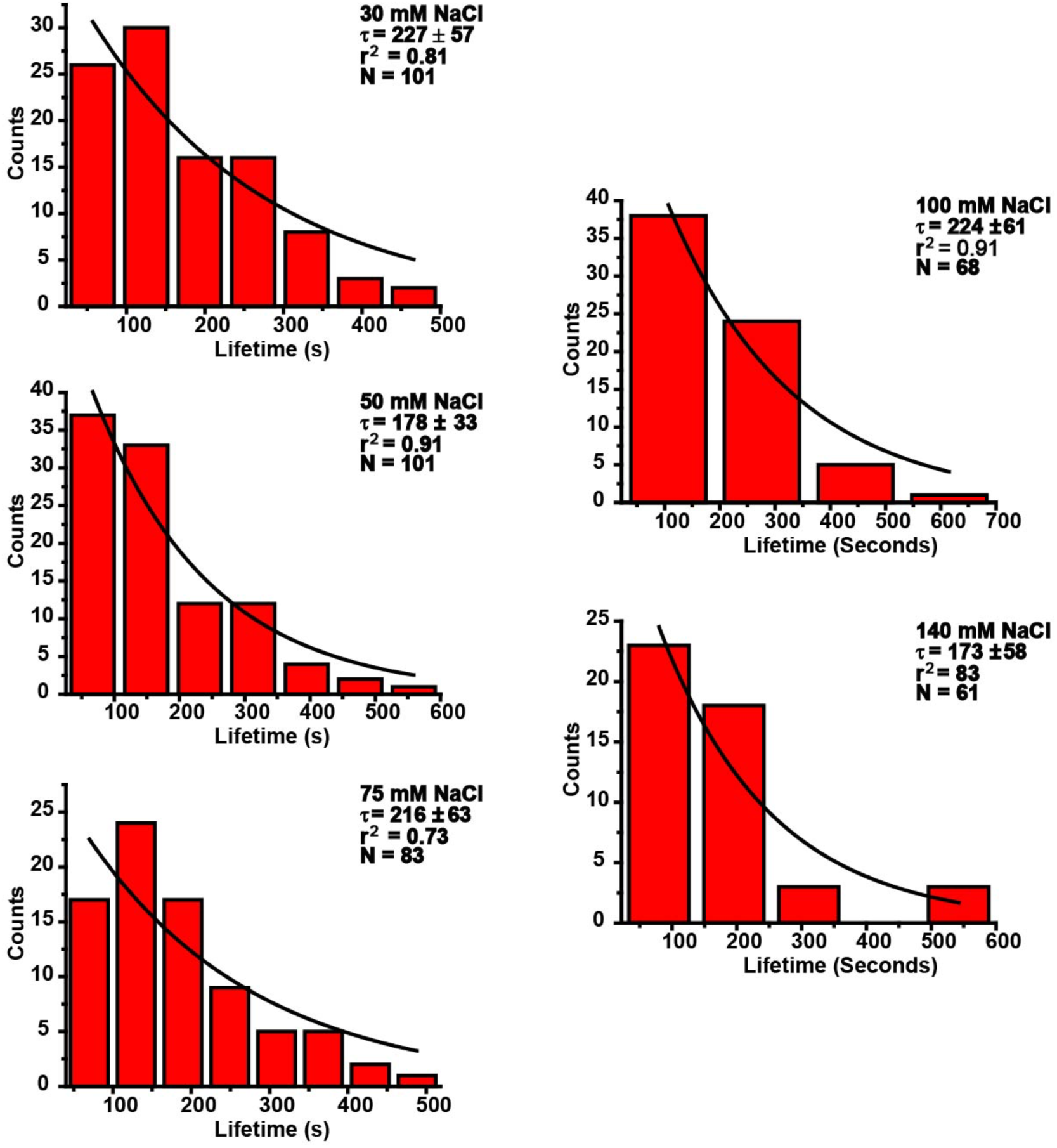
MutS Lifetime Calculations. Distribution of lifetime of EcMutS on a single mismatched DNA at various ionic strengths as indicated.

**Supporting Figure S5.**
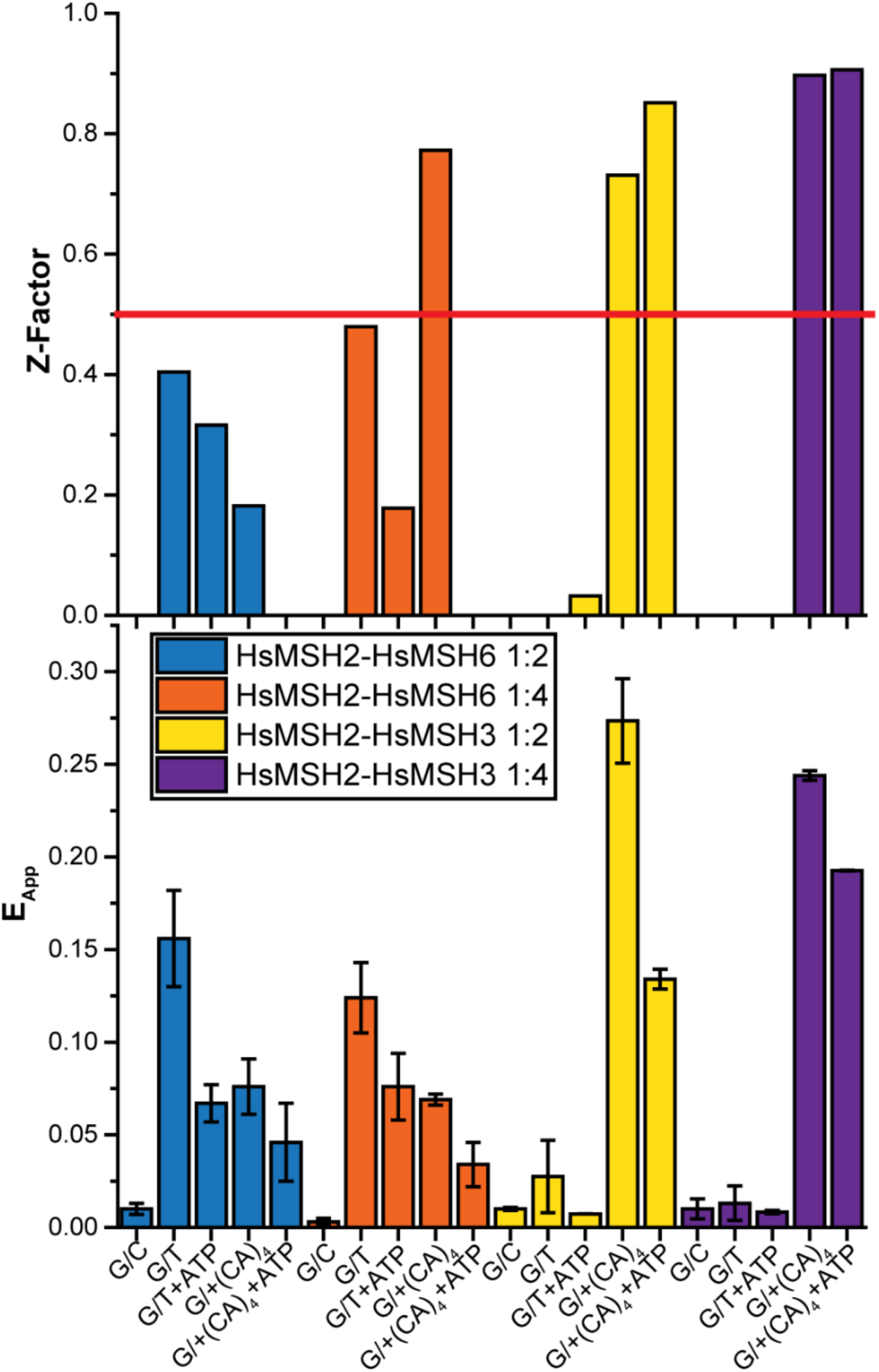
Z-Factor. (Top) Z-Factor quantification of assay effectiveness. Only values greater than zero shown. Red line shows 0.5 which indicates a high-quality drug screen. (Bottom) Composite E_App_ bar graph used to calculate Z-factor of 1:2 (HsMSH2-HsMSH6:DNA), 1:4 (HsMSH2-HsMSH6:DNA), 1:2 (HsMSH2-HsMSH3:DNA), 1:4 (HsMSH2-HsMSH3:DNA) for G/C; G/T; G/T + ATP;G/+(CA)_4_; G/+(CA)_4_ + ATP (mean ± SD).

## Notes

### Competing Interest Statement

The authors have declared no competing interest.

